# On the Effectiveness of Event-related Beta tACS on Episodic Memory Formation and Motor Cortex Excitability

**DOI:** 10.1101/078964

**Authors:** Verena Braun, Rodika Sokoliuk, Simon Hanslmayr

**Affiliations:** University of Birmingham (School of Psychology)

**Keywords:** beta oscillations;, episodic memory;, motor cortex excitability;, transcranial alternating current stimulation (tACS);

## Abstract

**Background:** Transcranial alternating current stimulation (tACS) is widely used to entrain or modulate brain oscillations in order to investigate causal relationships between oscillations and cognition.

**Objective:** In a series of experiments we here addressed the question of whether event-related, transient tACS in the beta frequency range can be used to entrain beta oscillations in two different domains: episodic memory formation and motor cortex excitability.

**Methods:** In experiments 1 and 2, 72 healthy human participants engaged in an incidental encoding task of verbal and non-verbal material while receiving tACS to the left and right inferior frontal gyrus (IFG) at 6.8Hz, 10.7Hz, 18.5Hz, 30Hz, 48Hz and sham stimulation for 2s during stimulus presentation.

In experiment 3, tACS was administered to M1 at the individual motor beta frequency of eight subjects. We investigated the relationship between the size of TMS induced MEPs and tACS phase.

**Results:** Beta tACS did not affect memory performance compared to sham stimulation in experiments 1 and 2. Likewise, in experiment 3, MEP size was not modulated by the tACS phase.

**Conclusions:** Our findings suggest that event-related, transient tACS in the beta frequency range cannot be used to modulate the formation of episodic memories or motor cortex excitability. These null-results question the effectiveness of event-related tACS to entrain beta oscillations and modulate cognition.

## Introduction

Brain oscillations represent regular fluctuations in the local field potential and play a crucial role in establishing synchronous firing patterns [1]. Especially oscillations in the beta frequency range (~13-30Hz) have been linked to a variety of cognitive and sensorimotor processes [2–6]. Beta power decreases, for example, have been shown to predict successful memory encoding [7,8]. Such desynchronized activity occurs in highly task relevant regions [9], and is negatively correlated with blood oxygenation level dependent (BOLD) activity [10]. It has further been demonstrated that power and phase of beta oscillations over the motor cortex influences the amplitude of TMS evoked potentials (MEPs) [11,12].

Despite the numerous associations between these processes and beta oscillations, the causal relationship between them remains unclear. Transcranial alternating current stimulation (tACS), an increasingly popular non-invasive human brain stimulation technique [13], has been suggested to provide this causal link between brain oscillatory activity and cognitive processes. Recent findings suggest that tACS entrains brain oscillations in a frequency specific way [14,15]. This modulation of underlying oscillatory activity can affect behaviour [16–19], interacts with underlying oscillatory activity [20–24], and elicits frequency specific neuronal spiking [25].

tACS could be a very beneficial and powerful method for cognitive research, if it was also able to modulate brain oscillations in a time-critical way [26]. During cognitive tasks brain oscillations show a very dynamic behaviour and are modulated in the range of seconds. However, most studies having demonstrated effects of tACS on behaviour applied tACS throughout cognitive tasks in a sustained way, resulting in stimulation durations of up to 20min [15–18,20,21,27]. This makes it difficult to directly link oscillatory activity associated with specific cognitive dynamics with the results of these tACS studies. In order to demonstrate that tACS is indeed a useful tool for modulating dynamic cognitive processes, tACS should be administered within a short period of time at certain phases of a cognitive task in event-related, randomized designs.

In the present study we sought to investigate the effectiveness of event-related beta tACS. In a series of experiments we explored whether tACS in the beta frequency range is effective in modulating two different processes: the formation of episodic memories and motor cortex excitability.

### Beta power decreases and episodic memory formation

Successful memory formation for verbal material has been associated with power decreases in the beta frequency range [8]. This beta desynchronization can be localized to the left inferior frontal gyrus (IFG) [10], a region which has been linked to successful semantic memory encoding in numerous studies [28]. Using rhythmic transcranial magnetic stimulation (rTMS), Hanslmayr et al [29] demonstrated a causal link between beta desynchronization in the left IFG and memory encoding. By artificially synchronizing the left IFG via rTMS in the beta frequency range, memory formation for words was impaired at beta but not at other frequencies. These findings provide a first causal link between beta power decreases and episodic memory. Experiments 1 and 2 aimed to replicate and extend these findings and further examine whether tACS may be a useful addition to TMS. In these two experiments, the differential and specific effects of beta tACS to the left IFG on the encoding of verbal material and the effects of beta tACS to the right IFG on the encoding of non-verbal material were investigated. Several studies report material-specific lateralization during episodic memory encoding with left frontal involvement during the encoding of verbal material and right frontal activation for non-verbal material [28,30,31]. Therefore beta (18.5Hz) tACS should only affect memory performance for words when administered to the left IFG (as has been shown by Hanslmayr et al. [29]) while right IFG stimulation should result in decreased memory performance for non-verbal material only.

### Beta phase and motor cortex excitability

Several simultaneous tACS-TMS studies investigated the causal relationship between beta power and corticospinal excitability [23,32], with a recent study investigating whether the phase of 20Hz tACS can modulate MEP amplitude [33]. However, in this study tACS was applied for 200s and a phase-dependent modulation was found for the last three MEPs only. In experiment 3 we aim to investigate whether 10s of tACS tuned to the individual motor beta frequency can entrain beta oscillations in the primary motor cortex [34] and hence lead to a modulation of the amplitude of TMS evoked MEPs according to the tACS phase.

Being able to modify beta oscillations within a short period of time in an event-related, randomized manner is crucial for quantifying the effectiveness of beta tACS and ultimately its usefulness for modulating dynamic cognitive processes. Although numerous studies have shown that tACS affects behaviour, the present three experiments could not find clear entrainment effects when applying tACS in the beta frequency range for shorter durations (2s, 10s). Indeed, Bayesian statistical analyses indicate that beta tACS has no effect compared to sham stimulation.

## Experiments 1 and 2

### Material and Methods

#### Participants

Participants were screened for contraindications against transcranial alternating current stimulation prior to the experiment [35].

36 subjects participated in experiment 1 (24 female; mean age: 20.03 +/- 2.38 years) and 36 in experiment 2 (24 female, mean age: 20.97 +/- 2.22 years).

All participants were right handed, had normal or corrected-to-normal vision and reported no history of neurological disease or brain injury. Informed consent was acquired from each subject prior to the experiment. All were naive to the hypotheses of the study and were fully debriefed at the end of the experiment. The study was approved by the ethics committee of the University of Birmingham.

### Stimulus Material

Word stimuli consisted of 270 nouns derived from the MRC Psycholinguistic Database, Version 2.00 [36]. These were divided into 18 lists of 15 words and were matched for word frequency, word length, number of letters, number of syllables, concreteness, and imaginability. Face stimuli consisted of 270 faces drawn from several face databases. The faces were emotionally neutral and were presented in black and white on a black background. These were divided into 18 lists of 15 stimuli and were matched for gender, hair colour, and approximate age. Stimuli were presented in a randomized order, and counterbalanced across subjects. 360 stimuli (180 words, 180 faces) were presented during encoding and retrieval, serving as old items in the retrieval period, 180 stimuli (90 words, 90 faces) were presented during retrieval only, serving as new items (figure 1).

### Experimental Setup and Procedure

Participants were seated approximately 80cm from a 19 inch LCD monitor (resolution: 1280 X 1024 pixels, 60Hz frame rate). Stimuli were presented in black on a grey background on the centre of the screen using the Psychophysics Toolbox extension for Matlab [37].

Before the start of the main experiment participants were familiarized with tACS and desensitized to the stimulation intensity in order to avoid adverse reactions.

### Encoding

During encoding, participants had to perform a pleasantness rating of a presented stimulus on a 4-point rating scale (very pleasant – very unpleasant). Answers were given manually by using the middle and index finger of both hands (figure 1A). During the 2s stimulus presentation, tACS was administered to the left and right IFG at different frequencies. In order to replicate and extend the findings from Hanslmayr et al. [29], tACS was applied at 18.5Hz. Furthermore two frequencies below and two frequencies above this frequency were chosen as control frequencies (stimulation frequencies: 6.8Hz, 10.7Hz, 18.5Hz, 30Hz, 48Hz, as well as sham stimulation). The sequence of the stimulation conditions was counterbalanced across subjects and pseudo randomized so that the same frequency and the same stimulation site did not occur in more than four consecutive trials.

**Figure 1.**
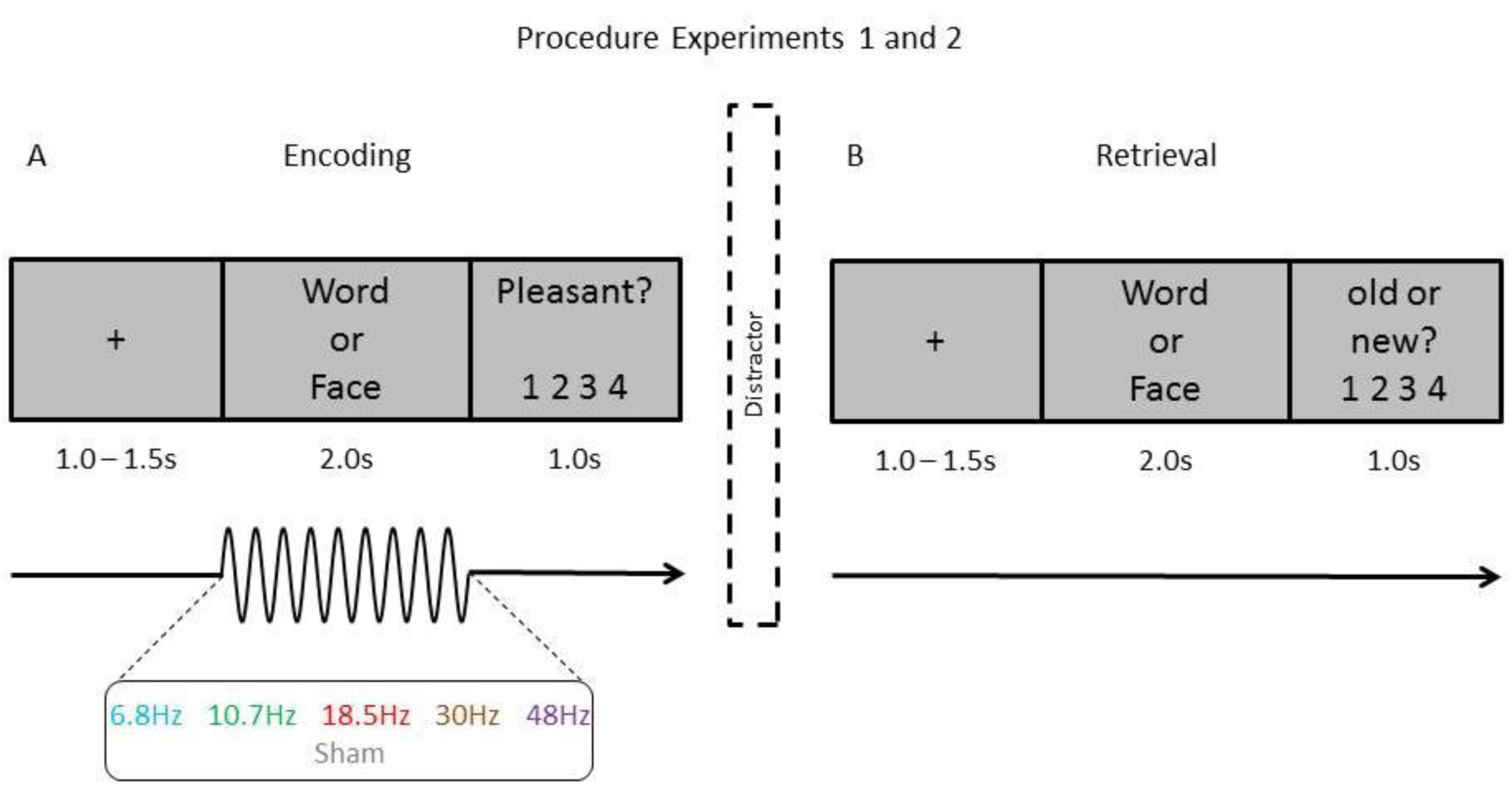
The experimental design for experiments 1 and 2 is shown. 360 stimuli (180 words, 180 faces) were presented during the encoding block (A). Participants had to rate the pleasantness of a stimulus on a 4-point rating scale *(very pleasant – very unpleasant).* During the 2s stimulus presentation, tACS was administered to the left and right IFG at 6.8Hz, 10.7Hz, 18.5Hz, 30Hz, 48Hz, as well as sham stimulation. The material was counterbalanced across subjects so that every stimulus was paired with every stimulation condition equally often throughout the experiment. During the retrieval block (B) the 360 stimuli presented during encoding as well as 180 new items (90 words, 90 faces) were shown. Subjects were asked to rate their confidence of an item being old or new on a 4-point rating scale (*very sure old* - *very sure new)*.

### Retrieval

Following the encoding section, two distractor tasks were used to ensure that the participants did not rehearse the study material. First, the participants were required to count aloud backwards in steps of 7 from a 3-digit number for 1min, after which time they were asked to rate the intensity of the stimulation induced sensations and phosphenes for every stimulation condition separately (see supplementary figure 3). These tasks were followed by the recognition phase. Here, the 360 items presented during encoding, along with 180 new items were presented in a randomized sequence. Subjects were asked to rate how confident they were that in item was old or new on a 4-point rating scale ranging from *very sure old* to *very sure new* (figure 1B).

### Transcranial Alternating Current Stimulation

Transcranial alternating current stimulation was delivered via a 4 channel DC Stimulator MC (NeuroConn, www.neuroconn.de). In experiment 1, the stimulation was applied via four donut-shaped rubber electrodes with a diameter of 5cm (14 cm2, NeuroConn, Ilmenau, Germany) at an intensity of 1mA (2mA peak to peak). In experiment 2, the stimulation was applied using round rubber electrodes with a diameter of 3.7cm (10.75 cm2, NeuroConn, Ilmenau, Germany) at an intensity of 0.8mA (1.6mA peak to peak). The resulting current density in both experiments was approximately 0.07mA/cm2. The stimulation electrodes were placed at EEG electrode positions FP1, C5, FP2, C6 (figure 2). These positions were selected using a neuro-targeting software (Soterix Medical Inc, New York, USA) which uses a finite-element model of a template adult brain to model the current distribution in the brain. Stimulation sites were chosen to result in the highest target field intensity in the left inferior frontal gyrus (figure 2A). In order to keep the sensations equal between stimulation conditions, the placement for the right IFG stimulation was derived by mirroring the montage for the left IFG stimulation onto the right hemisphere (figure 2B). Impedances were kept below 5kOhm using Ten20 conductive paste (Weaver and Company, Aurora/Colorado). In both experiments, tACS was applied at 6.8Hz, 10.7Hz, 18.5Hz, 30Hz, and 48Hz for 2s at stimulus onset during encoding (see figure1). Additionally, sham stimulation was applied. The current was ramped up and down at the beginning and at the end of the 2s stimulation period for around 300ms in all of the five stimulation frequencies.

**Figure 2.**
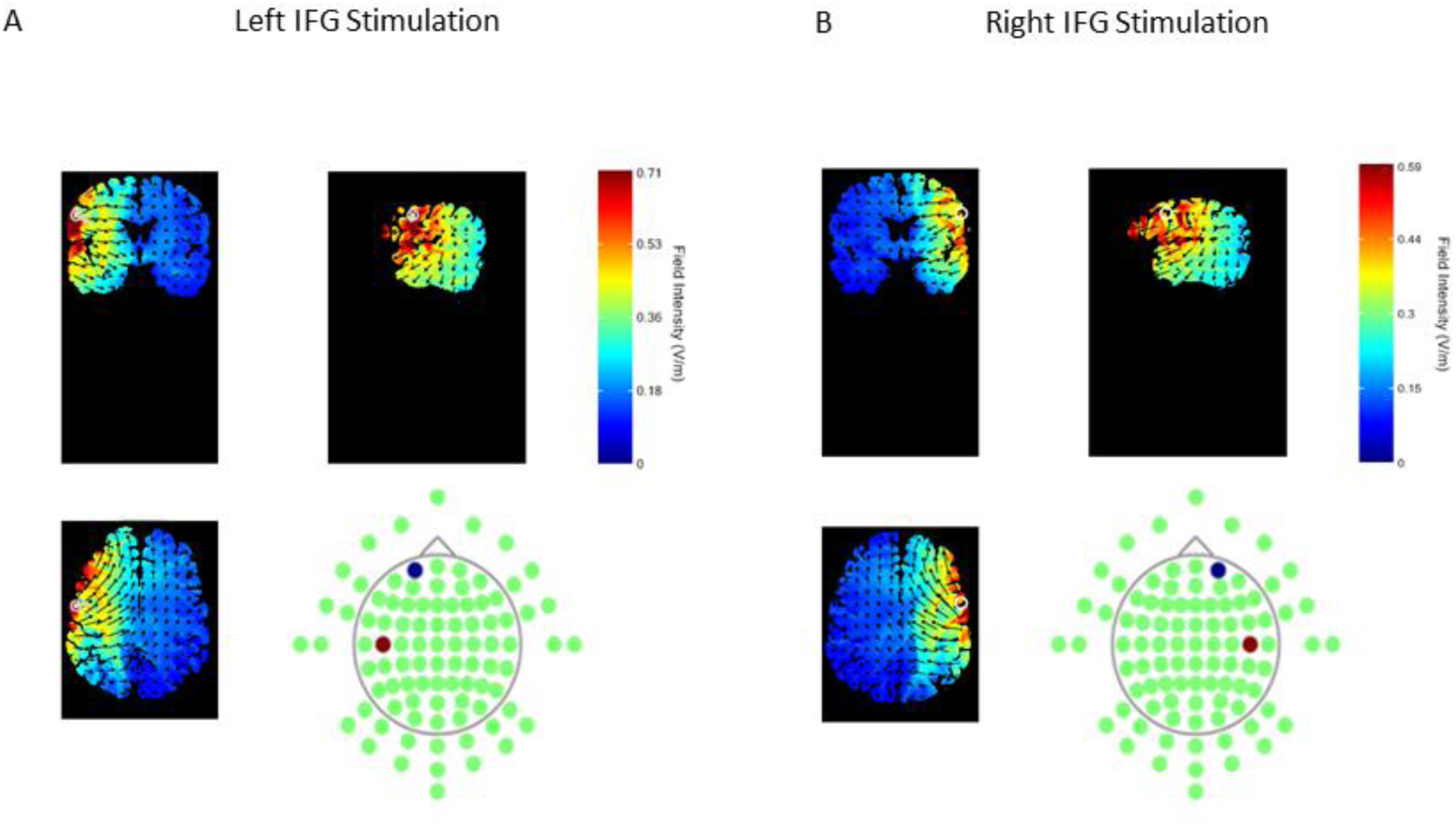
Stimulation electrode configurations for the left inferior frontal gyrus (A) and right inferior frontal gyrus (B). Optimal electrode placement for the left IFG (BA 9) was mirrored onto the right hemisphere. Current field intensity is shown using a finite-element model, provided by Soterix Medical Inc. The field intensities are shown for a stimulation of 2mA, whereas in the present experiments stimulation intensities of 1mA and 0.8mA were used.

### Data analysis

Correctly identified old stimuli (*hits*) were classified using a receiver operating characteristic (ROC) procedure. In order to control for individual response biases, every subject’s neutral response criterion was determined indicating which buttons a participant used for an old response and thus providing a bias free measure of memory strength [8].

The specific effects of left beta tACS on memory performance for verbal material and right beta tACS on memory performance for non-verbal material were assessed using Bayesian t-tests with a JSZ Prior (see supplementary material) [38]. The resulting Bayes factors (BF01) indicate how likely it is that the present data can be observed under the null hypothesis (there is no difference between the conditions) as compared to the alternative hypothesis (there is a difference between the conditions).

### Results

A 3-way ANOVA revealed no interaction between the stimulation frequency, stimulation site, and experiment for verbal material, F(5,350)=0.987, p=0.426. Hence, data of both experiments were merged into one dataset (figure 3A). No interaction between stimulation frequency and stimulation site could be observed, F(5,350) = 1.7, *p*=0.134. The difference between the left 18.5Hz stimulation and left sham stimulation was subjected to a Bayes t-test. The resulting Bayes factor indicates that the data is 7.564 times more likely to be observed under the null hypothesis, t(71)=0.204, p=0.839, BF01=7.564.

**Figure 3.**
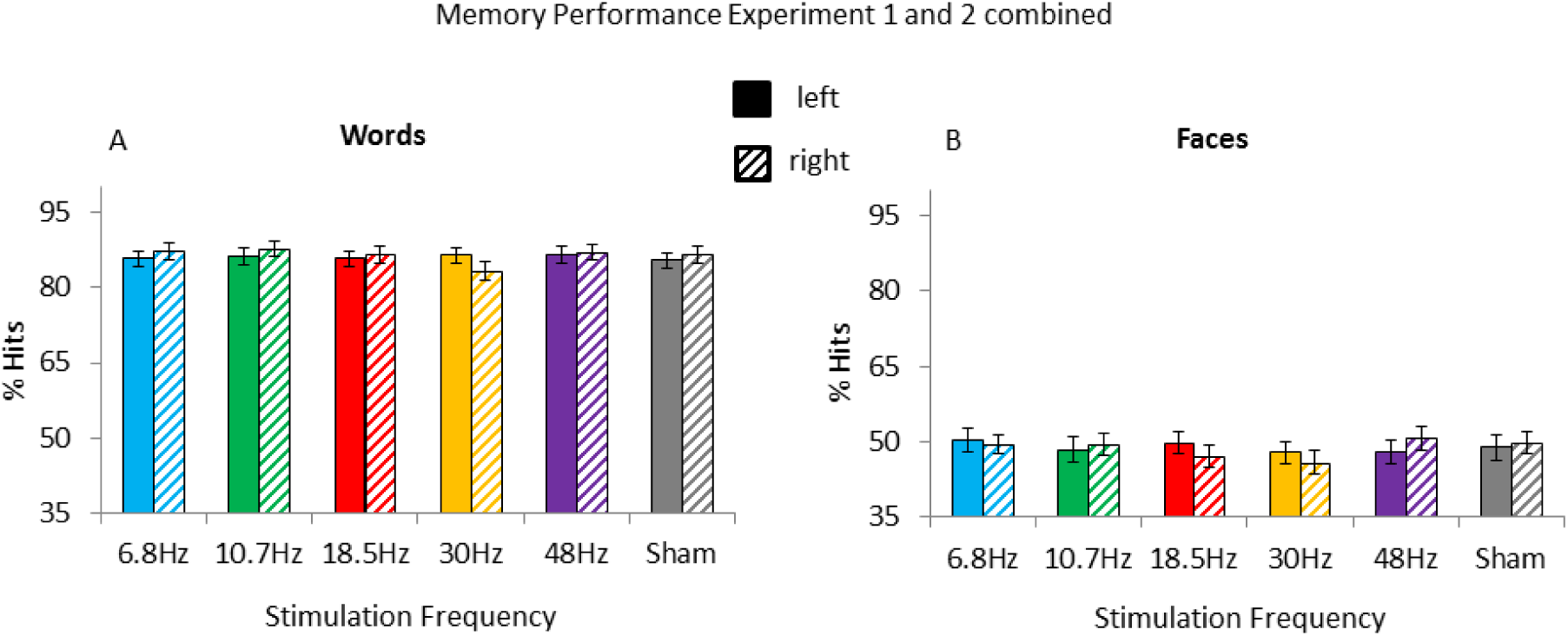
Memory performance for words (A) and faces (B) split by stimulation condition and stimulation site (data of experiments 1 and 2 combined). Bayesian t-tests indicate no difference in memory performance between left beta stimulation and left sham stimulation for words (A) and between right beta stimulation and right sham stimulation for faces (B). Error bars show standard errors of the mean.

A 3-way ANOVA for the non-verbal material revealed no interaction between stimulation frequency, stimulation site, and experiment F(5,350)=0.992, *p*=0.423. Therefore, data of both experiments were also combined (figure 3B). No interaction between stimulation frequency and stimulation site could be found, F(5,350)= 1.11; *p*=0.355. The difference between right 18.5Hz stimulation and right sham stimulation was subjected to a Bayes t-test, indicating that the data is 3.13 times more likely to be observed under the null hypothesis, t(71)=-1.377, *p*=0.173, BF01=3.13.

Memory performance for both experiments separately is depicted in supplementary figures 1 and 2. D’ values for sham stimulation are shown in supplementary table 1.

### Discussion experiments 1 and 2

In the present two experiments, we aimed to examine whether event-related, randomized tACS can be used to modulate beta oscillations during episodic memory formation. We specifically expected 18.5Hz tACS to decrease memory performance only for words when administered to the left IFG while right 18.5Hz stimulation should have resulted in decreased memory performance for non-verbal material only. Although similar protocols using TMS were able to show that left beta stimulation impairs encoding of verbal material [29], the present study failed to show such an effect. Beta tACS did not modulate the formation of episodic memories when applied in a temporally sensitive, event-related, randomized manner. These results indicate that, at this point, tACS in the beta frequency is not a suitable alternative to TMS. As tACS affects neurons in a more subtle fashion than TMS [39], tACS might not be strong enough to interfere with underlying oscillatory activity in such a short period of time [40].

## Experiment 3

### Material and Methods

#### Participants

Eight participants completed the experiment (all male; mean age: 29.375 +- 4.93 years).

Participants were screened for contraindications against tACS and TMS prior to the experiment [35,41]. All participants were right handed, had normal or corrected to-normal vision and reported no history of neurological disease or brain injury. Informed consent was acquired from each subject prior to the experiment. The study was approved by the ethics committee of the University of Birmingham.

#### Experimental Setup and Procedure

Due to safety considerations, the experiment was split into two consecutive sessions. Both sessions consisted of the same experimental procedure.

#### Transcranial Magnetic Simulation

Transcranial magnetic stimulation (TMS) was delivered with a Magstim Rapid stimulator via a 70mm double coil (magstim; www.magstim.com) to the left motor cortex at 110% motor threshold (identified without active tACS, but over the tACS electrode) every 2.5s – 4.5s [12], resulting in 420 TMS pulses per session. The stimulation site (M1) was defined as the position on the scalp that elicited the strongest MEP response.

#### Transcranial Alternating Current Stimulation

TACS was delivered via a 4 channel DC Stimulator MC (NeuroConn, www.neuroconn.de). The stimulation was applied using round rubber electrodes with a diameter of 3.7cm (10.75 cm2, NeuroConn, Ilmenau, Germany) at an intensity of 0.7mA (1.4mA peak to peak), resulting in a current density of 0.065 mA/cm2. Two electrode montages were used in order to investigate the efficiency of montages with one electrode directly over the target area [23,32] as compared to montages with the target area in between the stimulation electrodes (as used in the previous tACS memory experiments presented). This resulted in stimulation electrodes being placed at M1, and EEG electrode positions Pz and Fp1. This setup allowed us to use the same reference electrode, Pz, in both stimulation conditions, so that a randomized stimulation protocol could be used. Impedances were kept below 5kOhm using Ten20 conductive paste (Weaver and Company, Aurora/Colorado).

#### tACS-TMS Procedure

During the experiment, participants were seated comfortably in front of a computer screen. No task was involved. Subjects were instructed to keep their hands as relaxed as possible, while looking at a fixation cross in the middle of the screen. Single pulse TMS was delivered throughout the experiment over the tACS electrode placed at M1 (figure 4), while participants received tACS at their individual motor beta frequency (see supplementary material). tACS was applied for 10s, followed by a 10s period without stimulation. The electrode montage with which the stimulation was delivered (i.e. FP1-Pz or M1-Pz), was pseudorandomised so that the same montage was never repeated more than twice. Motor evoked potentials were measured from the first dorsal interosseous (FDI) muscle of the right hand using Ag-AgCl EEG electrodes (BrainAmp MR plus, Brainvision). Every 17-18 trials, participants were given a slight break and the TMS coil was cooled down, resulting in four tACS-TMS blocks per session. The tACS artefact was recorded from one Ag-AgCl EEG electrode placed at Cz, referenced to the right mastoid.

**Figure 4.**
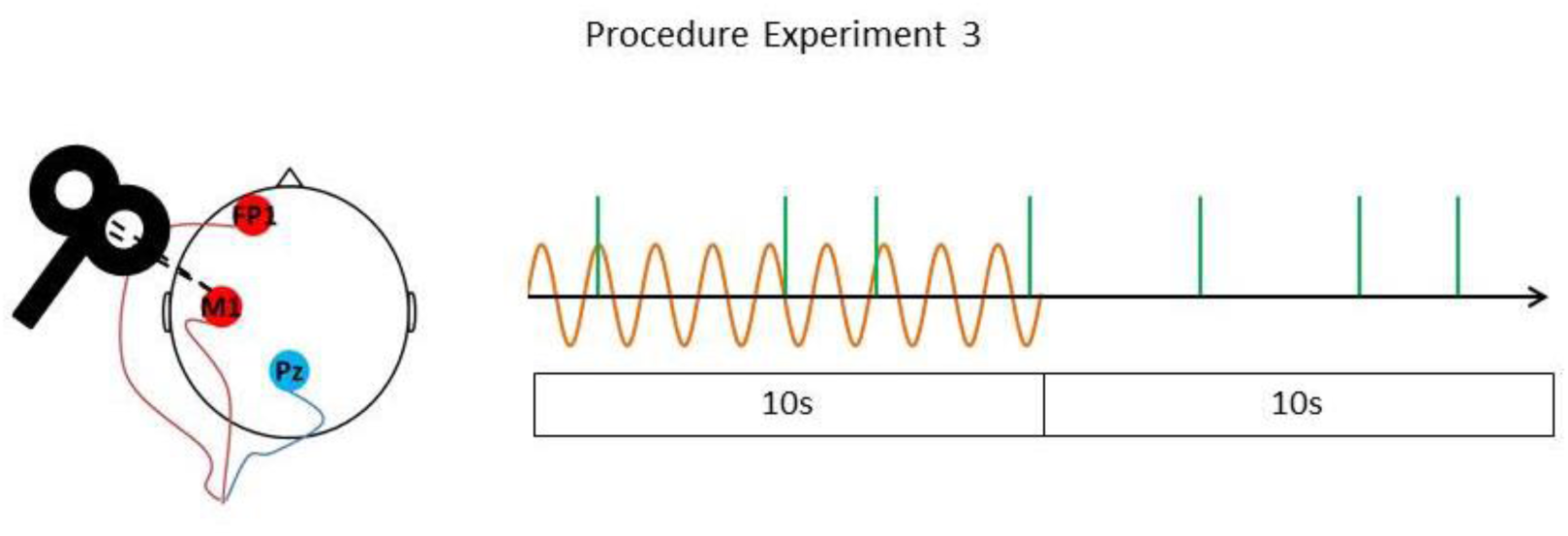
tACS-TMS procedure for experiment 3 is shown. TMS pulses (depicted in green) were delivered throughout the trial every 2.5s-4.5s to the left motor cortex over the tACS electrode placed at M1. tACS was applied at the individual motor beta frequency (depicted in orange) for 10s followed by a 10s period without tACS. 2 different tACS montages were used: M1-Pz and FP1-Pz. MEPs were measured from the first dorsal interosseous (FDI) muscle of the right hand. Each session consisted of 70 trials.

#### Data Analysis

Data were analysed using FieldTrip [42], the CircStat toolbox [43], and in-house MATLAB scripts. Due to noise in the MEP data produced by the tACS stimulation, amplitudes of the motor evoked potentials were not easily accessible. Therefore, MEP data were subjected to a time-frequency composition using Morlet wavelets and baseline corrected (baseline window: −0.15s to −0.05s). MEP amplitude was defined as the peak of the mean signal change between 20Hz and 50Hz 0-50ms following the TMS pulse, which revealed clear MEPs (see supplementary figure 4). In order to adjust for noise introduced by the breaks and possible changes in position of the TMS coil, MEP amplitudes in every block were z-transformed, ensuring that data from every block were comparable. To extract phase angles of tACS at the moment of each TMS pulse, EEG data recorded from electrode Cz were Hilbert transformed. In order to test for phase entrainment effects, circular to linear correlations between the normalised MEP amplitudes and tACS phase were calculated. Additionally, the data were binned into four different tACS phase bins centred around 0° (*peak)*, 90° (*falling flank*), 180° (*trough*), 270° (*rising flank*) [33], and normalised MEP amplitudes at those tACS phase bins were subjected to a repeated measures ANOVA.

#### Results

Overall, there was no significant difference in MEP size between tACS trials in either of the montages and no-tACS trials (figure 5A); Montage 1 (M1-Pz): t(7)=−0.951, *p*=0.373, BF01= 2.07; Montage 2 (FP1-Pz): t(7)=1.350, *p*=0.219 BF01=1.5. Normalised single trial MEP amplitude by tACS phase collapsed across both montages is shown in figure 5B. No correlation between MEP amplitude and tACS phase could be found; overall: _cl_ = 0.0131, *p*=0.7732; Montage 1: _cl_ = 0.0195, *p*=0.7548 (supplementary figure 5A); Montage 2: _cl_ = 0.0420, *p*=0.2675 (supplementary figure 5B).

**Figure 5.**
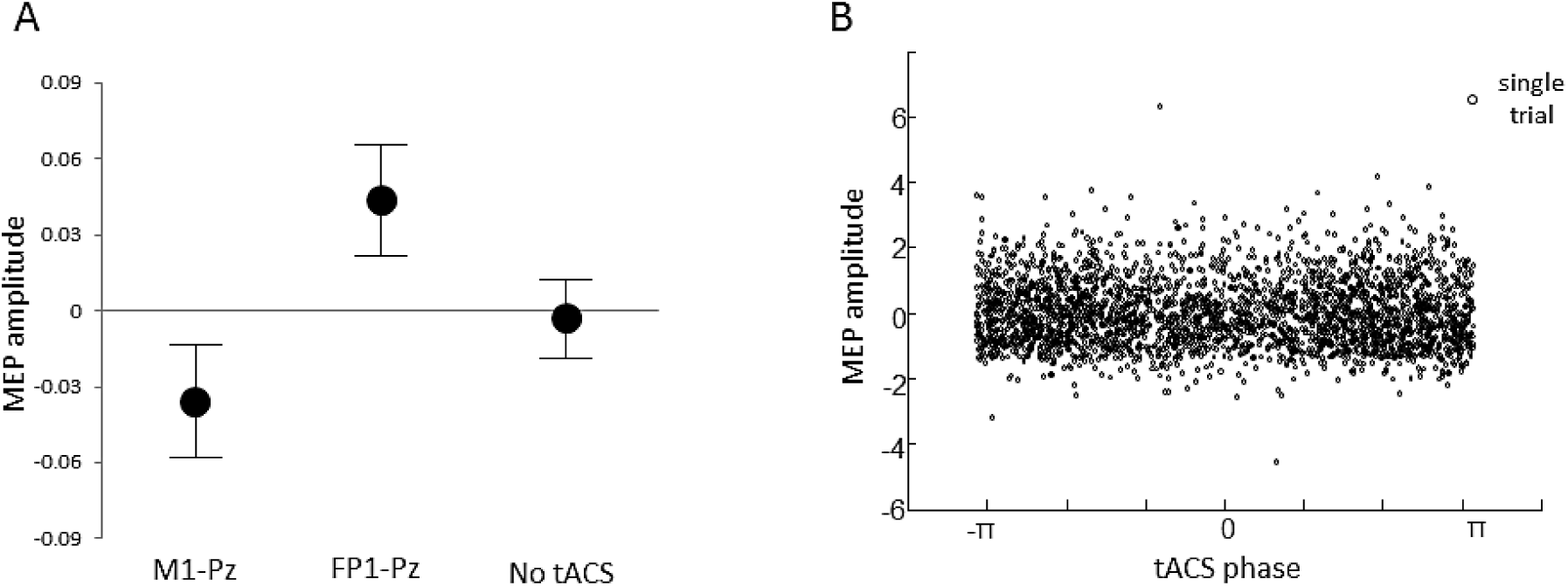
(A) Normalised mean MEP amplitude split by tACS condition. Error bars show standard errors of the mean. (B) Single trial MEP size by tACS phase.

tACS phase at the moment of the TMS pulse did not show an effect on MEP size (figure 6), F(3, 21) = 0.473, *p*=0.704. No interaction between the phase bins and tACS montage could be found either, F(3,21)=0.847, *p*=0.484, however, there was a main effect for tACS montage with higher MEP amplitudes at M2 (FP1-Pz) than M1 (M1-Pz); F(1,7)=7.902, p=0.026 (see supplementary material).

Individual motor beta frequencies and resting motor thresholds are depicted in supplementary table 2.

**Figure 6.**
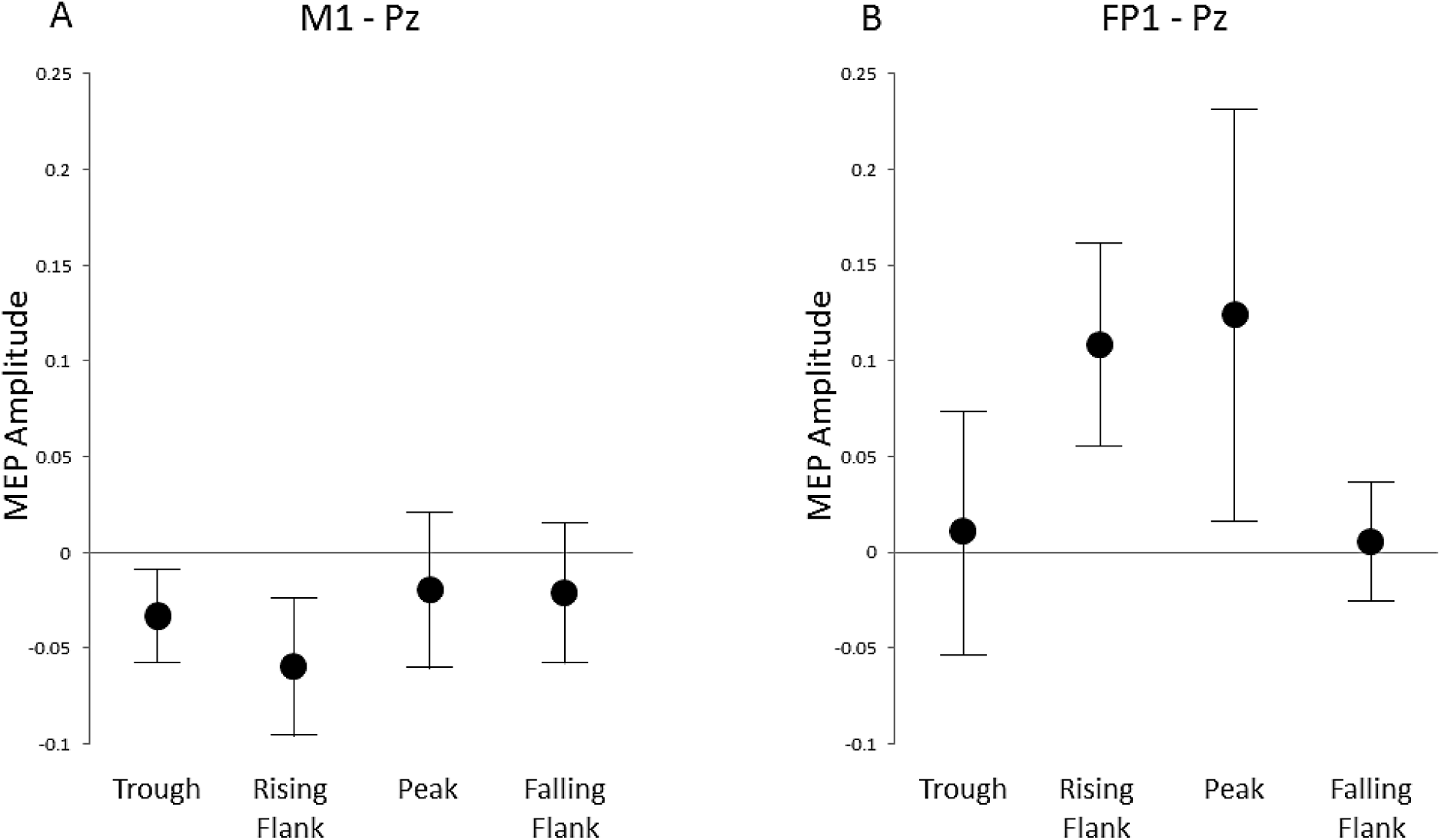
Mean normalised MEP amplitude split by 4 different tACS phase bins: 180° (trough), 270° (rising flank), 0° (peak), 90° (falling flank). A. tACS Montage 1: M1-Pz. B. tACS Montage 2: FP1-Pz. Error bars show standard errors of the mean.

#### Discussion experiment 3

The aim of experiment 3 was to examine if it is possible to entrain beta oscillations in the primary motor cortex using tACS. By applying tACS to M1 at the individual motor beta frequency, we investigated the relationship between TMS induced MEPs and tACS phase. As in the first two experiments and as previous studies indicate [33], we did not find a clear entrainment effect; the MEP size was not modulated by the tACS phase. We believe that this is due to the rather short tACS stimulation period (10s) we used, since other studies that reported an effect of beta tACS stimulation on general MEP size used longer stimulation periods [23,32].

### Conclusion

During cognitive tasks brain oscillations, and beta oscillations in particular, show a highly dynamic behaviour. For instance beta oscillations decrease in power within a couple of milliseconds during memory processing or movement execution followed by a subsequent increase in amplitude. If tACS may be useful in answering causal questions about these dynamics, our findings would need to show that tACS at beta is capable of modulating oscillatory behaviour in a similar time range. However, in the series of experiments reported here we found that event-related, randomized, transient tACS in the beta frequency range does not modulate the formation of episodic memories or motor cortex excitability.

This failure to find an effect of beta tACS on cognition or cortical excitability—and the finding that such statistically positive effects are unlikely—reveals that the effectiveness of tACS is a complex issue. On the one hand, tACS applied over minutes appears to be effective in modulating behaviour and brain oscillations [15–18,20–22,33]. And indeed, if tACS is used to synchronize and desynchronize distant brain regions, shorter stimulation durations (1s- 1.8s) in other frequencies seem to be successful [26]. On the other hand, the present findings indicate that tACS applied in the range of seconds in order to modulate brain oscillations in one brain area is not effective [40,44]. Before being able to utilize beta tACS to draw conclusions about the causal relationship between oscillatory brain activity and cognitive processes, several issues regarding the use of beta tACS protocols need to be addressed, such as current distribution in the brain, optimal electrode placement, recommended stimulation intensities, recommended stimulation durations etc. Though a growing body of modelling studies addresses these issues [45–49], the respective models have to be validated extensively by experimental data, before it will be possible to apply tACS more effectively in cognitive research [50]. Event-related beta tACS could then be a useful and promising method. Yet as long as these problems remain unsolved, tACS may remain ineffective in unravelling the causal relationship between transient beta oscillatory activity and cognitive function.

## Acknowledgments

This work was supported by grants awarded to S.H. by Deutsche Forschungsgemeinschaft [Emmy Noether Programme Grant HA 5622/1-1]; and the European Research Council [Consolidator Grant Agreement 647954].

